# Bootstrap aggregating improves the generalizability of Connectome Predictive Modelling

**DOI:** 10.1101/2020.07.08.193664

**Authors:** David O’Connor, Evelyn M.R. Lake, Dustin Scheinost, R. Todd Constable

## Abstract

It is a long-standing goal of neuroimaging to produce reliable generalized models of brain behavior relationships. More recently data driven predicative models have become popular. Overfitting is a common problem with statistical models, which impedes model generalization. Cross validation (CV) is often used to give more balanced estimates of performance. However, CV does not provide guidance on how best to apply the models generated out-of-sample. As a solution, this study proposes an ensemble learning method, in this case bootstrap aggregating, or bagging, encompassing both model parameter estimation and feature selection. Here we investigate the use of bagging when generating predictive models of fluid intelligence (fIQ) using functional connectivity (FC). We take advantage of two large openly available datasets, the Human Connectome Project (HCP), and the Philadelphia Neurodevelopmental Cohort (PNC). We generate bagged and non-bagged models of fIQ in the HCP. Over various test-train splits, these models are evaluated in sample, on left out HCP data, and out-of-sample, on PNC data. We find that in sample, a non-bagged model performs best, however out-of-sample the bagged models perform best. We also find that feature selection can vary substantially within-sample. A more considered approach to feature selection, alongside data driven predictive modeling, is needed to improve cross sample performance of FC based brain behavior models.

## Introduction

A long-standing goal of neuroimaging research has been to establish generalizable links between brain structure/function and behavior or traits. A general approach is to identify discriminating imaging features which when incorporated into a statistical model can reliably estimate an observable phenotype for novel participants. The ultimate aim is to derive clinically actionable diagnoses or intervention strategies from imaging data [1]. In order to achieve a clinically actionable model, both feature engineering and model building are important. However, a simple model may still perform well with robust features that have a large effect size, yet even the most complex model cannot perform well given poorly curated features (garbage in, garbage out). In neuroimaging, an ideal feature set is seldom seen due to a combination of factors including site effects, physiological noise, hardware noise, and, in particular, small sample sizes, not to mention the complexity of the underlying neuronal activity itself. In order for the combination of a feature set and model to be clinically actionable in the real world these issues, which are ubiquitous, need to be overcome. In other words, in virtually all cases the model is beholden to features.

Functional magnetic resonance imaging (fMRI) based functional connectivity (FC) is a commonly used feature set for developing brain imaging-based models of observable phenotypes. Though initial investigations using fMRI focused on functional specialization of brain regions, it has been shown that widespread neural processes can contribute to higher level brain function. This implies that functional integration, rather than discrete specialization, is likely the key to characterizing more complex phenotypes [2], [3]. As a result, FC based models commonly use the functional connectome, a map of the functional connections between all pairs of brain regions (or nodes, defined by an atlas), as the starting point for feature selection [4]–[7]. This framework for analyzing brain function mostly incorporates undirected, first order, pairwise estimates of connectivity between brain regions [8]. Even still, dependent upon the size of the node atlas, these approaches typically yield upwards of 30,000 candidate features. These massive, and often disparate, sets of features are both hard to interpret, and easily allow for overfitting, particularly when working with small samples.

Approaches to avoiding overfitting depend upon the goals of the analysis, and the type of model being built; whether it be a predictive or explanatory model. These strategies are not mutually exclusive [9], but can be contrasted as being biased towards either model performance (predictive) or requiring there be a causal link between features and the observed phenotype (explanatory). Here, we will focus on the former. Recently there has been much interest in predictive modeling approaches that estimate brain-behavior relationships using network theory, with some success [7], [10]–[14]. These strategies, are geared toward model performance-based assessments, and attempt to avoid overfitting using cross validation (CV) [15]. However, as is the case with predictive frameworks in general, outcomes are agnostic to the consistency of model parameters or selected features. Predictive approaches do not guarantee a model will translate to other datasets. Instead, they ensure model performance is robust to the diversity within the dataset used to generate the model (training data). Notably, within-sample performance estimates can vary, especially when datasets are small (<100 participants) [16]. This is not to say that models built within one dataset using CV to avoid overfitting will always fail to generalize. Some studies have shown good performance out-of-sample [17]–[20]. For example, in each study it was a requirement that a given feature be selected in a minimum number of CV folds before being included in the external application of the model. This approach begins to resemble an ensemble learning method.

Ensemble learning refers to a group of statistical methods which act to combine multiple models, in order to boost performance [21]. Ensemble learning can be used to alleviate overfitting. An example ensemble learning method is bootstrap aggregating, or bagging [15], [22]. Bagging is based on training models in multiple subsets of a training data set, or bootstraps, and aggregating model parameters and features across subsets into one feature set and one model. The CV models cited above, which have been applied successfully out-of-sample, have in essence used a bagging approach. Bagging has been applied to fMRI data before to determine brain state [23], and for brain parcellation (to define an atlas) [24], and has also been shown to boost within-sample performance of resting state FC based brain-behavior models [25]. Here we apply a bagged predictive modeling approach, using connectome predictive modeling (CPM) as the model framework. We develop a brain-behavior models of working memory FC and fluid intelligence (fIQ) using the typical approach of using only one feature selection and model training step, and using an alternative bagging approach, combining information from multiple iterations of feature selection and model formation. We then show, using out-of-sample testing, that bagging improves model generalizability. Our results provide a framework for building more translatable brain behavior models.

## Methods

### Datasets

Data from the Human Connectome Project (HCP), specifically the S900 sample, [26] and the Philadelphia Neurodevelopmental Cohort (PNC), N=1000 [27], were used. From these datasets, N=827 participants from the HCP, ages 21-35, and N=788 participants from the PNC, ages 8-21, were included based on the availability of preprocessed T_1_-weighted images, working memory fMRI scans, and fIQ measures.

For the HCP dataset, fIQ was measured using a 24-item version of the Penn Progressive Matrices assessment, scores ranged from 4 to 24, with a mean of 16.78 ± 4.7 [28]. In the PNC dataset, the 24- and 18-item versions of the Penn Matrix Reasoning Test were used [29], [30], scores ranged from 0 to 23, with a mean of 11.85 ± 4.06. In both datasets, the score corresponds to the number of correct responses.

For the HCP, MRI data were acquired on a 3T Siemens Skyra. The fMRI scans were collected using a slice-accelerated, multiband, gradient-echo, echo planar imaging (EPI) sequence (TR = 720ms, TE = 33.1ms, flip angle = 52°, resolution = 2.0mm^3^, multiband factor = 8, left-right phase encoding, scan duration = 5:01). The T_1_-weighted structural scans were collected using a Magnetization Prepared Rapid Gradient Echo (MPRAGE) sequence (TR = 2400ms, TE = 2.14ms, TI = 1000ms, resolution = 0.7mm^3^) [31]. For the PNC, the MRI data were acquired on a 3T Siemens TIM Trio. fMRI scans were collected using a multi-slice, gradient-echo EPI sequence (TR = 3000ms, TE = 32ms, flip angle = 90°, resolution = 3mm^3^, scan duration = 11:39). T_1_-weighted structural scans were collected using an MPRAGE sequence (TR = 1820ms, TE = 3.5ms, TI = 1100ms, resolution = 0.9375×0.9375×1mm) [32].

### Preprocessing

For the HCP, the HCP minimal preprocessing pipeline was used on these data [33], which includes artifact removal, motion correction, and registration to MNI space. All subsequent preprocessing was performed in BioImage Suite [34] and included standard preprocessing procedures [10], including removal of motion-related components of the signal; regression of mean time courses in white matter, cerebrospinal fluid, and gray matter; removal of the linear trend; and low-pass filtering.

For the PNC, structural scans were skull stripped using an optimized version of the FMRIB’s Software Library (FSL) pipeline [35]. Slice time and motion correction were performed in SPM8 [36]. The remainder of image preprocessing was performed in BioImage Suite [34] and included linear and nonlinear registration to the MNI template; regression of mean time courses in white matter, cerebrospinal fluid, and gray matter; and low-pass filtering.

FC matrices were generated from the working memory fMRI data using the Shen atlas [37], and Pearson’s R as the distance metric. CPM was performed as in [38], with the FC matrices as the explanatory variable, and fIQ as the target variable, with one exception; partial correlation was used at the feature selection step [39]. At this point, motion (mean frame to frame displacement) was regressed from both the predictor and target variable prior to correlation, before feature selection. Data analysis was performed primarily in python, with one exception; the signed rank statistical test for comparing model performances which was performed in MATLAB. Figures were generated in python using matplotlib [40] and seaborn [41], with the flowchart generated using BioRender.

### Modeling Protocol

Overall, we perform 20 test/train splits to assess the impact of participant inclusion in the training set on model performance. The following protocol explains what happens within each split. The protocol is also shown in Figure 1.

**Figure 1.**
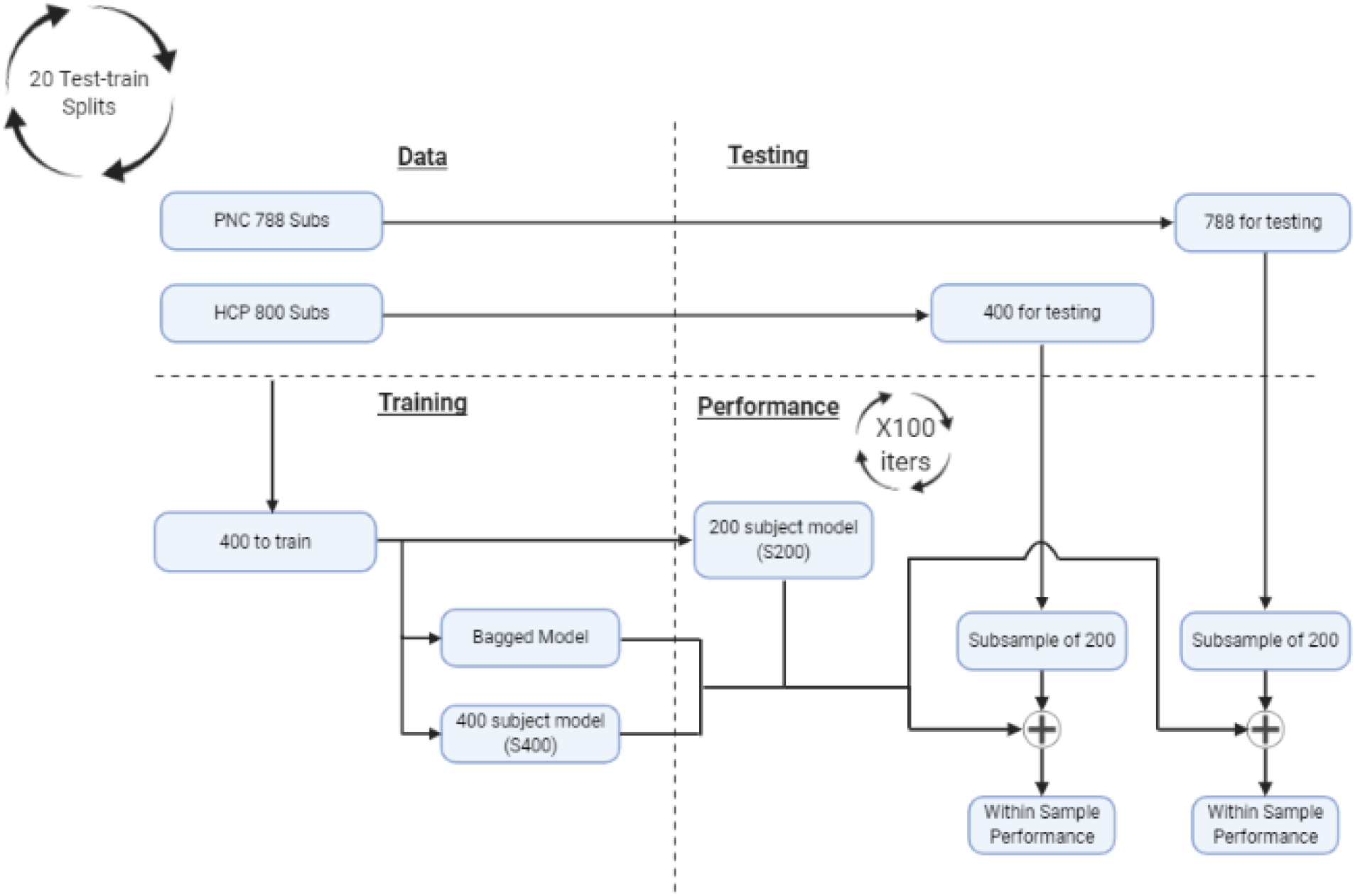
Description of the analytic workflow. The PNC data is only used for out-of-sample testing. The HCP data is split into train and test samples. The train sample is used to train 3 types of models: (1) a bagged model, and a non-bagged models based on (2) the whole train set (400 participants), and (3) 100 non-bagged models based on random subsamples of the train set (200 participants in each case). All models are then tested within-sample on the test HCP sample, and out-of-sample on the PNC dataset.

#### Model Training

800 subjects are randomly selected from the N=827 HCP subjects, and equally divided into a train and test sample. The test sample is used exclusively to test within-sample performance, and none of the participants in the train sample are included in testing, see Fig 1. The train sample is used to build three types of models: one bagged (1) and two non-bagged (2 & 3).

1. To build the bagged model, all 400 participants in the train set are used. 100 iterations of feature selection and model building are performed on bootstraps of 200 randomly selected participants. The model parameters, the slope and intercept, and features are aggregated to form one bagged model. The slope and intercept of the bagged model are the mean across bootstraps. A mean feature vector is formed, which is the frequency with which each feature (edge) is selected across all bootstraps. Both feature set and model parameters together will be termed the ‘bagged’ model.
2. The first non-bagged model is generated using all 400 participants without any bagging, this will be termed the ‘S400’ model.
3. The second non-bagged model is comprised 100 models generated from the train sample without bagging, each based on a randomly selected 200 participants, these will be termed the ‘S200’ models.

Given the 20 test-train splits, our full protocol yields 20 bagged models, 20 S400 models trained on 400 subjects, and 2000 S200 models trained on 200 subjects.

#### Model Testing

The HCP test sample is used to assess the within-sample performance of all models, and the PNC data is used to assess out-of-sample performance. In both within-sample, and out-of-sample testing, each of the model types (bagged, S400, and S200) are tested on 100 random subsamples of 200 participants from the within and out-of-sample test sets. I.e. 100 random subsamples of 200 participants are generated from the HCP test set. For each of these subsamples, the bagged model is evaluated, one S200 model is evaluated, and the S400 model is evaluated. This framework yields 100 measures of performance for each model within-sample. The same subsampling of the PNC is done to evaluate out-of-sample performance to yield 100 measures of out-of-sample performance.

The bagged model is first tested including every feature which occurred at least once in any one bootstrap of the model training. Following this, the feature vector is iteratively thresholded. First, to include features which occur in 10% or more bootstraps, then 20% or more, rising in 10% increments, to those that occur in 90% or more of bootstraps. For each threshold, the bagged models’ performance is assessed within and out-of-sample, as described above.

Model performance is quantified as the strength of the correlation between the predicted and actual fIQ. Differences in model performance are assessed using the signed Wilcoxon rank test, as the model performance scores were not Gaussian.

## Results

The performance of non-bagged and bagged models (with all features included) in predicting fIQ, within and out-of-sample are shown in Figure 2. Performance metrics are separated based on the 20 test-train splits. Performance of all models vary depending on the test train split, but generally the S400 performs best in sample, and the bagged model performs best out-of-sample.

**Figure 2.**
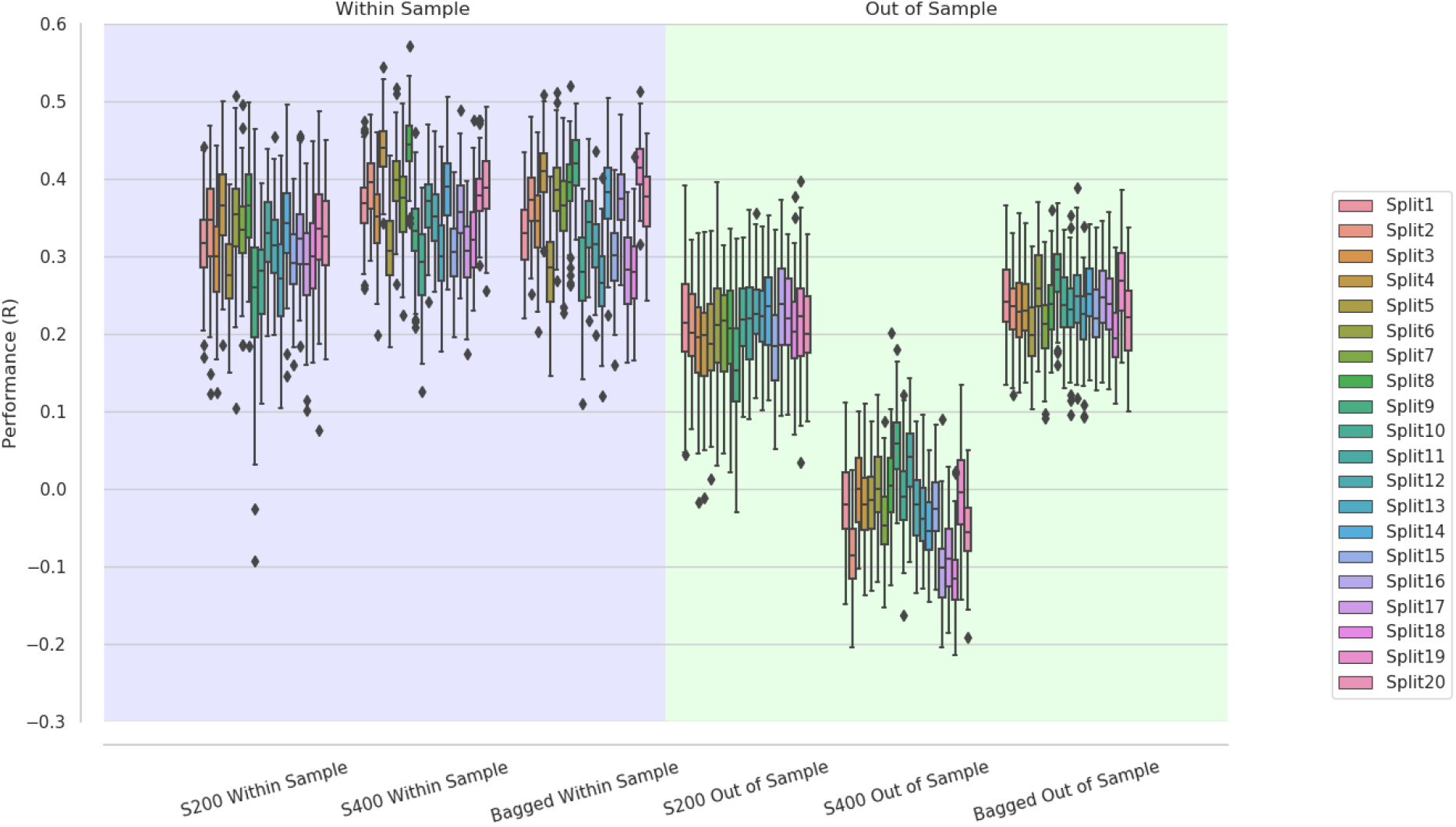
The first three columns (shaded in purple) show performance of models within-sample, color-coded (in rainbow) by train/test split. Single models are trained on either 200 (S200, column one) or 400 (S400, column two) subjects from the HCP training set. The bagged model (column three) is trained on 100 bootstraps of 200 subjects from the HCP train sample. All models are tested within-sample on 100 random subsamples of 200 subjects from the HCP test set. Columns four, five and six (shaded in green) show the performance of the same models out-of-sample using random subsamples of 200 subjects from the PNC data.

In Figure 3, performance metric distributions are shown without differentiating by test-train split. Within-sample, the S400 models perform best, better than the bagged models (W = 1,515,316, p = 2.18e-88), which perform better than the S200 models (W = 1,544,707, p = 1.53e-98). Out-of-sample, the bagged models perform best, better than the S200 models (W = 1,508,226, p = 5.06e-86), which performed better than the S400 models (W = 2,000,283, p = 0).

**Figure 3.**
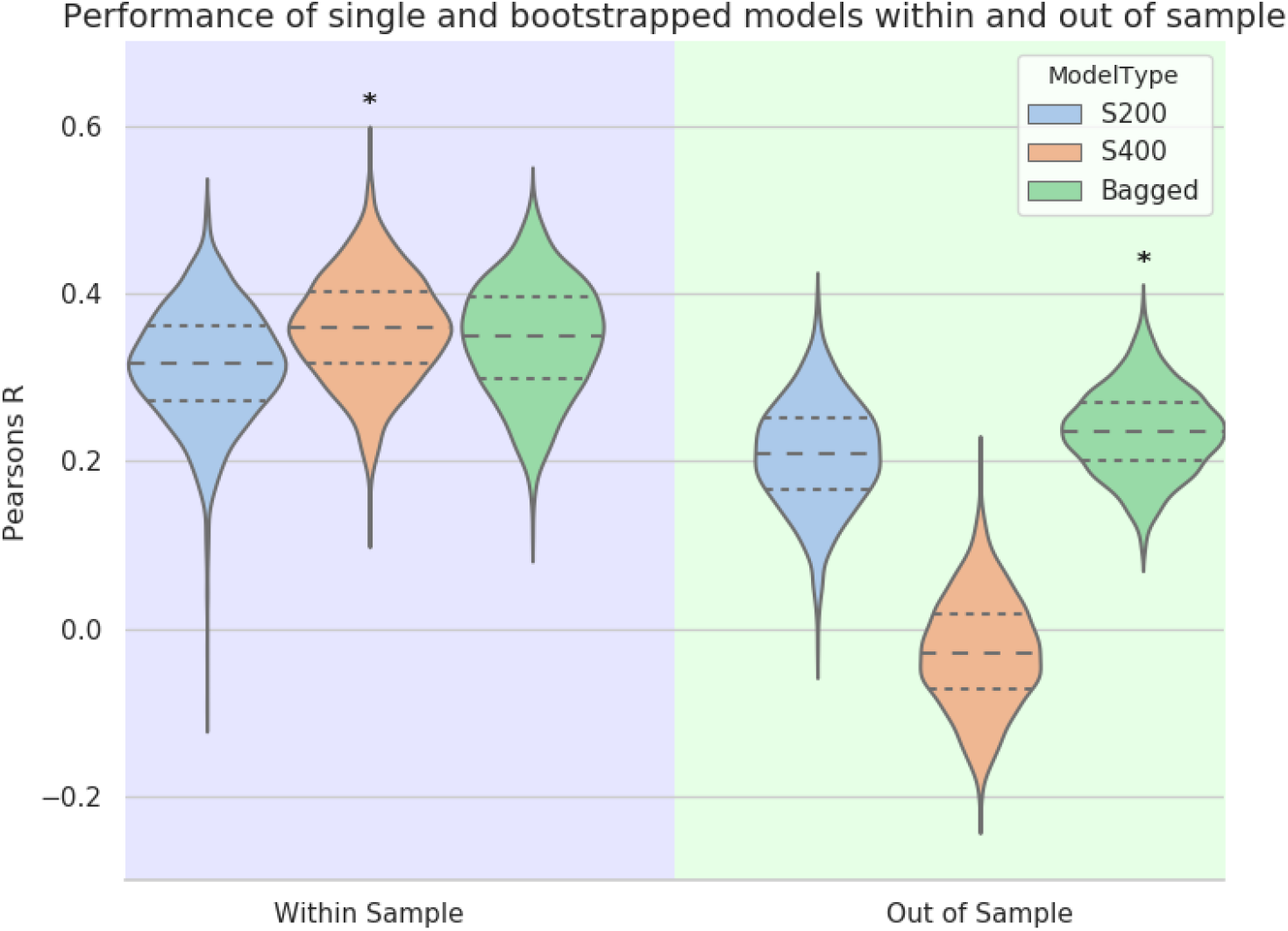
Column one (shaded in purple) shows performance within-sample. Non-bagged models are trained on either 200 or 400 subjects from the HCP training set, (S200/S400). The bagged model is trained on 100 bootstraps of 200 subjects from the HCP training set. All models are then tested on 100 random subsamples of 200 subjects from the HCP test set. The second column (shaded in green) shows the performance of the same models tested on random subsamples of 200 subjects from the PNC data. Star designates the best performing models within-sample (S400) and out-of-sample (Bagged) based on Wilcoxon signed rank test.

The distribution of feature occurrence in the bagged models is shown in Figure 4. On average 47.5% ± 3.3%, or 17,008 ± 1193 out of a possible 35,778 unique connections, are selected at least once across bootstraps using our undirected FC matrix (which contains 268 nodes). On average 29 ± 24 features occur in >=90% of bootstraps. With CPM, edges (features) are selected based on how their strength (the correlation of the signal between two nodes) correlates with the target variable across individuals within the training set. This is a statistically thresholded step. When creating the bagged models, the R values at this feature selection step are indicative of whether a feature occurs frequently across bootstraps, as shown in Figure 5. An example from building one bagged model is shown in Figure 5. The R values ranged from −0.306 to 0.31.

**Figure 4.**
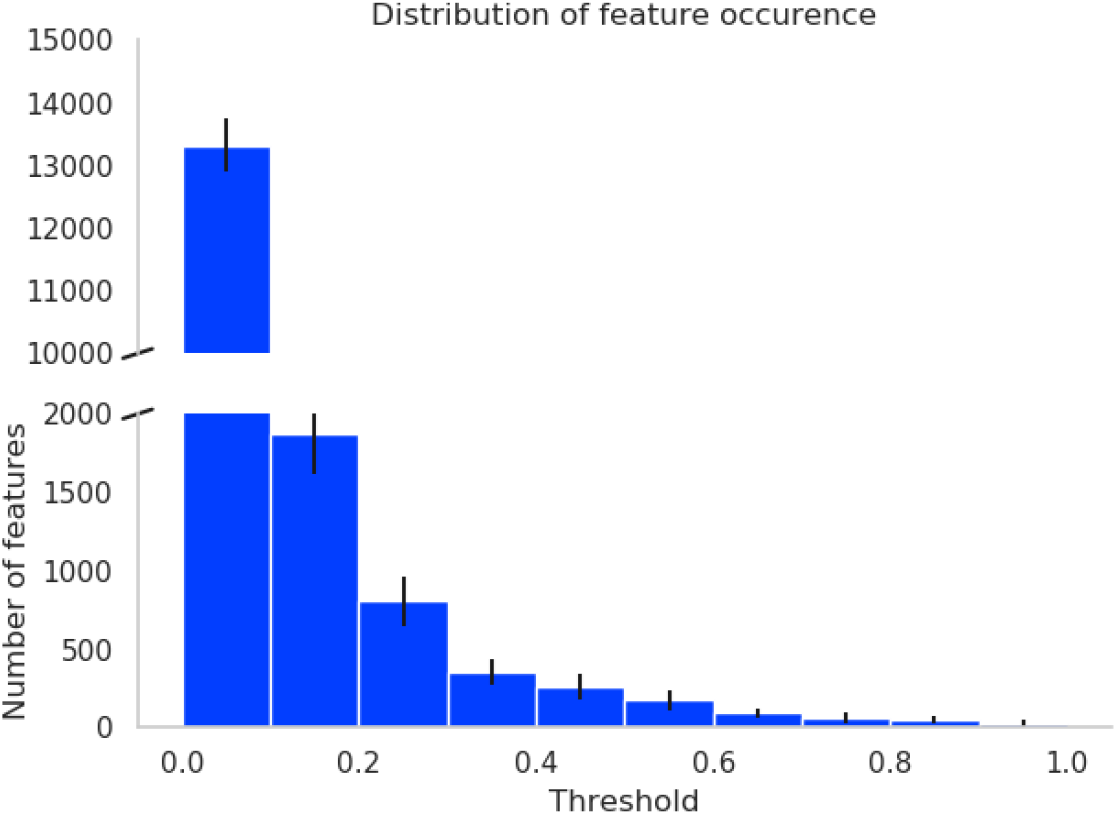
Distribution of feature (edge) occurrence across bootstraps for the bagged models. On average, nearly 14,000 features only occur in between 0% and 10% of bootstraps.

**Figure 5.**
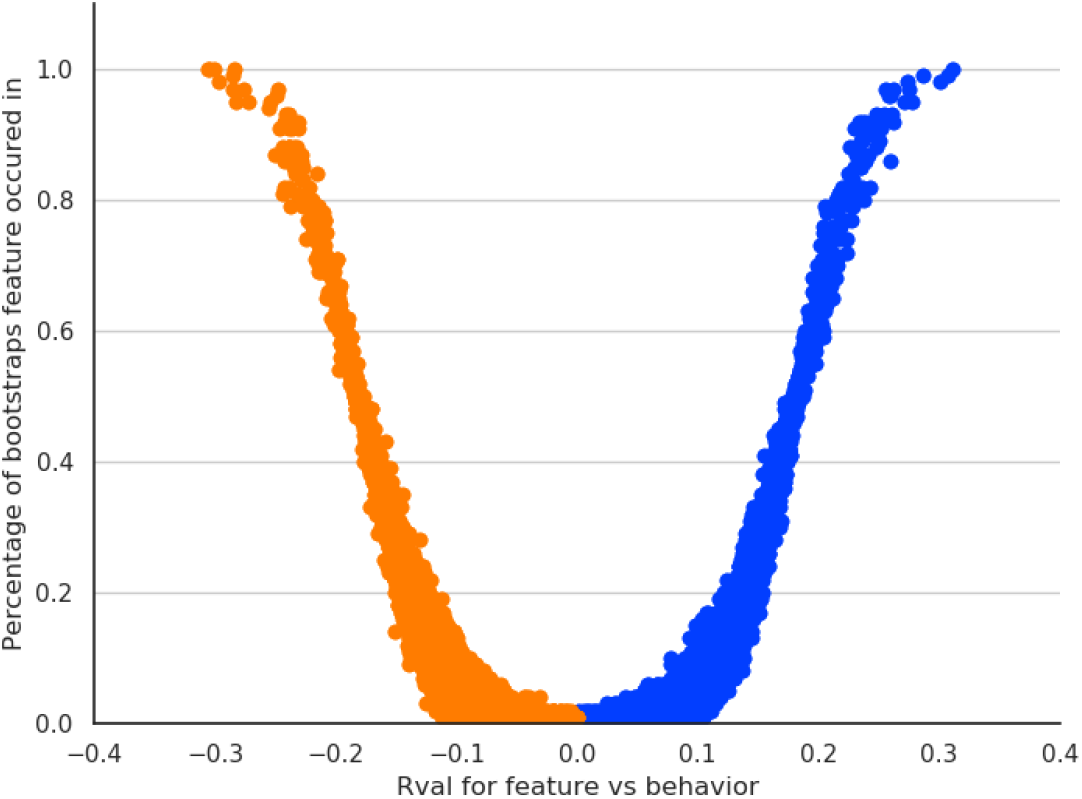
Feature occurrence relative to its correlation with the behavioral measure determined at the feature selection step, as determined in the feature selection when creating a bagged model. The blue dots represent those features positively correlated with the target variable, while the orange dots are the negatively correlated features.

The effect of thresholding the feature vector on the bagged model’s performance is assessed within-, Figure 6, upper panel, and out-of-sample, Figure 6, lower panel. Within- and out-of-sample, bagged model performance did not change with feature thresholding up to 50%. Beyond this, out-of-sample performance fell as feature threshold increased (fewer features included).

**Figure 6.**
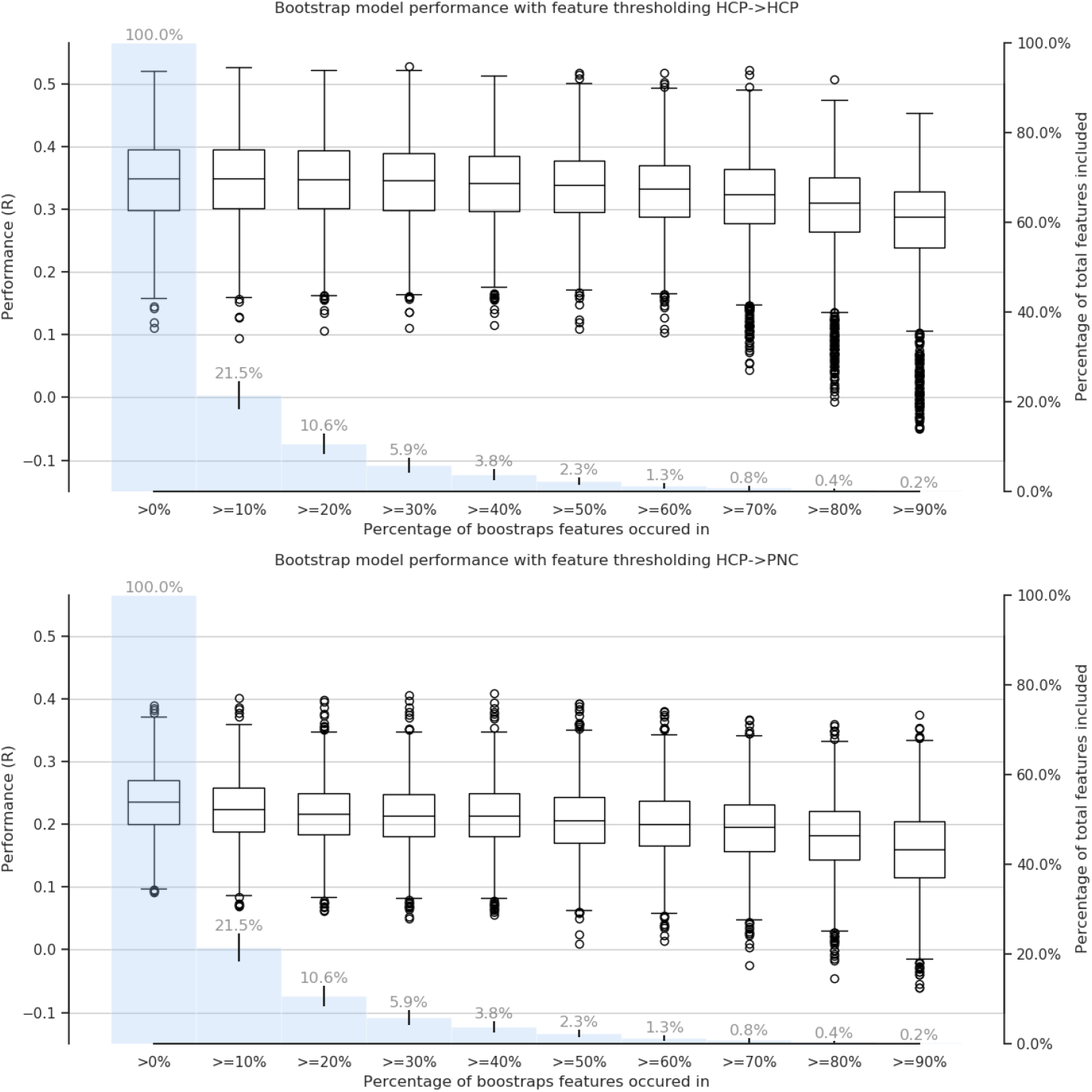
A. shows the performance of the HCP bagged models within-sample, as the feature threshold is increased (reducing the number of features included). B. shows the performance of the HCP bagged models out-of-sample, as the feature threshold is increased. In the background of both plots, a histogram of the percent of features included is shown in blue.

Out-of-sample bagged model performance remains significantly higher than S400 model performance even when the bagged model features are thresholded at 90% occurrence (p =1.34e-320, W = 1989241). On average, in the bagged models, only 29 ± 24 (mean ± standard deviation, SD) features meet this cut-off across splits. Across all S400 models, the average number of features is 1521 ± 324. This means that greater predictive power is obtained out-of-sample from roughly a fiftieth the number of features when an aggressively thresholded bagged model is compared to a non-bagged model which uses all individuals in the train set to build one model.

The out-of-sample performance of the bagged model remains significantly better than the S200 model up to a 40% feature reoccurrence threshold (p = 4.23e-04, W = 1091554), after which the S200 model performs better. At a 40% threshold the bagged model contains on average 257 ± 82 edges. The average number of edges in an S200 model is 896 ± 246. In other words, when the thresholded bagged model contains ~30% of the edges contained within the typical S200 model, predictive performance out-of-sample of these approaches is equivalent.

## Discussion

In this study we find that an ensemble method, in this case bagging, can improve the generalizability (out of sample performance) of brain-behavior models over standard cross validation. Three types of model are evaluated; bagged models, models trained on half the train set (S200), and models trained on the full train set (S400). Within-sample evaluation shows that S400 models perform best, followed by the bagged model, with S200 performing worst. Out-of-sample testing is used as a proxy for generalizability. Out-of-sample, the bagged models perform best, followed by S200, with the S400 models performing by far the worst. Generation of the bagged models requires estimating how frequently a given feature is selected across bootstraps. High variance is exhibited, with almost half of all features in the feature set being selected at least once. There is a strong relationship between effect size (R) and how many bootstraps a given feature is selected within. The effect of thresholding the feature set, based on feature occurrence, on the bagged models’ performance is evaluated. Within-sample, the bagged models’ performance remains steady until reaching a threshold of 50%. Out-of-sample the performance shows a slow decrease with increasing thresholding. Bagging or other ensemble methods are good alternatives to CV when generalizability is of key concern.

Our study is performed in response to a general increase in research which focuses on predictive modeling approaches that utilize FC measures to uncover meaningful brain behavior relationships. Ultimately, with these studies, a clinical application is desirable. However, present FC based predictive models are yet to perform well enough in terms of accuracy, sensitivity or specificity to have an impact on clinical practice. Generalizability is of particular importance in a clinical setting, and presently a major problem for predictive models in neuroimaging. In some cases, sophisticated models, which have shown much success in other fields, such as convolutional neural networks, have been applied to model brain-phenotype relationships [7], [42]. However, they are yet to provide a generalizable model with sufficiently high-performance fit for clinical application. Given the failure of highly flexible and complex models to fit these data in a generalizable manner, and the myriad of potential confounds that exist in raw neuroimaging data, models themselves are not the barrier to generalization. What may be holding us back is feature selection.

More care must be taken with respect to feature selection, and how the features relate to the target variable, which is arguably more of an explanatory approach. Explanatory approaches are usually contrasted with predicative approaches, however, the predictive-explanatory dichotomy isn’t necessary [9], one can take more care with ensuring the use of reproducible features, while still defining success based on prediction performance. This is possible using ensemble modeling methods which incorporate the feature selection step, as in this study. This allows for the derivation of a stable estimate of both model parameters and features within the predictive framework. These methods, allied to out-of-sample performance assessment, can provide a pathway to generalization. When done correctly, these approaches allow us to assess if a model reproducibly relates features to a phenotype we are interested in.

A common method for avoiding overfitting, and promoting generalization, is CV [43], [44]. Though there are exceptions [17]–[20], models using CV are more commonly built and evaluated in one relatively homogenous sample and sometimes subsequently tested out-of-sample. If we are to follow a similar approach, we should pick a model solely based on performance within-sample. By this token, we would choose our S400 model. It performs on average better within-sample than the bagged and S200 models. However, out-of-sample, it performs worst of the three models, with the bagged models performing best. This suggests that bagging is preferable to CV, though these are not mutually exclusive. CV is necessary to get a more balanced measure of performance but it does not guarantee replication. Studies which demonstrate successful replication that use CV have in-essence implemented a form of bagging through feature aggregation. For example, testing on an out-of-sample data-set using only the most commonly occurring features across CV folds [17]–[20]. However, in these implementations, the maximum number of feature selection iterations would be 10. Given that with the 100 bootstraps, we get almost 50% of features occurring at least once, and that another study by Wei et al also recommend ~100 bootstraps [25], bagging may provide an improvement on identifying noisy features compared to 10-fold cross validation, as well as giving more insight into feature variation. Furthermore, the greater the fraction of subjects from the training set that are included in the feature selection step, in the case of 10-fold CV it would include 90% of the subjects, the less likely the features are to vary across folds. This may seem like an advantage, but, depending on sample size, obtaining information on how smaller subsamples influence variance in feature selection may be beneficial. A potential downside of this approach is feature occurrence is a fairly simple way of uncovering within-sample variance. Hypothetically, it is possible that some features are highly related and are not adding unique information. This would indicate using dimensionality reduction [45], [46] and/or a regularized modeling approach [14]. However, it is possible to integrate these methods into an ensemble approach, and it would be prudent to assess the impact of subsampling on the parameters generated.

Despite increased performance out-of-sample, the bagged models did not reach their own level of within-sample performance. This may suggest that the bagged models are overfit to the training data, especially in light of the number of features included in the bagged model at the lower levels of feature thresholding. Overfitting is defined as “The production of an analysis which corresponds too closely or exactly to a particular set of data, and may therefore fail to fit additional data or predict future observations reliably” [47]. This is commonly thought to mean that the model has fit too much noise, or signal of non-interest. In the case of FC based models, overfitting can occur as a result of modeling signal of non-interest, or as a result of fitting a subset of the signal of interest too well. To simply ascribe the difference in performance to overfitting signal of non-interest, would be to ignore studies suggesting age and sex-based differences in FC [13], [48]–[51], along with a host of other factors. One of the strengths of ensemble learning is that it can incorporate variation within dataset through subsampling, but it can only do that to the extent to which the training data exhibits variance. This may explain why the bagged models’ performance decreased out of sample with feature thresholding. At a low threshold for feature inclusion, it is likely that we capture some features relevant to estimating fIQ in in the out-of-sample test set, particularly given nearly half of all possible features were included. Increasing the threshold for feature inclusion would equate to including features that are more relevant to the training sample and not the test sample. In this study, models are trained on HCP data, which is comprised of adults (ages 21-35), and tested on PNC data, which includes adolescents (ages 8-21). In addition to capturing noise specific to the training sample, our model is likely not capturing features that are specific to adolescent cognition that are undoubtedly present in the out-of-sample test set from the PNC. Another confounding factor is that the working memory tasks in the HCP and PNC data sets have non-negligible differences. The experimental designs of each sample use different visual stimuli, and working memory loads. Additionally, the target variables are derived from different, though related, assessments of fIQ. These factors likely contribute to the difference in performance we observe between within- and out-of-sample testing across models. Our results suggest there is a tradeoff between fitting the variance we are interested in (adult cognition), and providing a generalizable model of fIQ. Generalizability encompasses both model parameters and features that are neither too “noisy”, nor too biologically specific. In this respect, the bagged model seems to capture features that improve generalizability, but not sufficiently to match the level of within-sample performance. It is likely that overfitting and an imperfect match between samples negated perfect generalization.

Though we are in an era where predictive modeling is the focus of much research, we do not yet have a complete understanding of how the features we identify, in this case FC data obtained during a working memory task, vary across cohorts. How we should adapt our approaches to account for noise and sample-specific features is evolving. We assert that an ensemble approach to predictive modeling can help discern features that are more relevant to model generalizability.

## Conclusions

Bagging models allows for greater model performance and generalizability. The within-sample boost in performance, in light of including all features in performance assessment, would suggest overfitting. However, out-of-sample performance was also significantly better, suggesting better model generalizability with bagging. Bagging also provides an estimate of the stability of feature selection across variable training subsamples. Within-sample bagged models maintain performance when decreasing the number of features up to a point. This is not replicated out-of-sample, as the performance decreases with removal of features.

## Data and code availability statement

The HCP data that support the findings of this study are publicly available on the ConnectomeDB database (https://db.humanconnectome.org). The PNC data that support the findings of this study are publicly available on the database of Genotypes and Phenotypes (dbGaP, accession code phs000607.v1.p1); a data access request must be approved to protect the confidentiality of participants. Code for conducting the analyses described here can be found at https://github.com/YaleMRRC/CPMBaggingAnalysis

## Acknowledgements

Data were provided in part by the Human Connectome Project, WU-Minn Consortium (Principal Investigators: David Van Essen and Kamil Ugurbil; 1U54MH091657) funded by the 16 NIH Institutes and Centers that support the NIH Blueprint for Neuroscience Research; and by the McDonnell Center for Systems Neuroscience at Washington University. The remainder of the data used in this study were provided by the Philadelphia Neurodevelopmental Cohort (Principal Investigators: Hakon Hakonarson and Raquel Gur; phs000607.v1.p1). Support for the collection of the data sets was provided by grant RC2MH089983 awarded to Raquel Gur and RC2MH089924 awarded to Hakon Hakonarson. All subjects were recruited through the Center for Applied Genomics at The Children’s Hospital in Philadelphia.

## References

[1] T. Insel et al., “Research Domain Criteria (RDoC): Toward a,” Am. J. Psychiatry Online, no. July, pp. 748–751, 2010.

[2] N. B. Turk-Browne, “Functional interactions as big data in the human brain,” Science, vol. 342, no. 6158. American Association for the Advancement of Science, pp. 580–584, 2013.

[3] J. Rissman, A. Gazzaley, M. D’esposito, and H. H. Wheeler, “Measuring functional connectivity during distinct stages of a cognitive task.”

[4] F. X. Castellanos, A. Di Martino, R. C. Craddock, A. D. Mehta, and M. P. Milham, “Clinical applications of the functional connectome,” Neuroimage, vol. 80, pp. 527–540, Oct. 2013.

[5] K. Dadi et al., “Benchmarking functional connectome-based predictive models for resting-state fMRI,” Neuroimage, vol. 192, pp. 115–134, May 2019.

[6] M. R. Arbabshirani, S. Plis, J. Sui, and V. D. Calhoun, “Single subject prediction of brain disorders in neuroimaging: Promises and pitfalls,” Neuroimage, vol. 145, pp. 137–165, Jan. 2017.

[7] U. Pervaiz, D. Vidaurre, M. W. Woolrich, and S. M. Smith, “Optimising network modelling methods for fMRI,” Neuroimage, vol. 211, p. 116604, May 2020.

[8] O. Sporns, “Network attributes for segregation and integration in the human brain,” Current Opinion in Neurobiology, vol. 23, no. 2. pp. 162–171, Apr-2013.

[9] M. D. Rosenberg, B. J. Casey, and A. J. Holmes, “Prediction complements explanation in understanding the developing brain,” Nat. Commun., vol. 9, no. 1, p. 589, Dec. 2018.

[10] E. S. Finn et al., “Functional connectome fingerprinting: identifying individuals using patterns of brain connectivity,” Nat. Neurosci., vol. 18, no. 11, pp. 1664–1671, Nov. 2015.

[11] S. M. Smith et al., “A positive-negative mode of population covariation links brain connectivity, demographics and behavior,” Nat. Neurosci., vol. 18, no. 11, pp. 1565–1567, Nov. 2015.

[12] M. D. Rosenberg, E. S. Finn, D. Scheinost, R. T. Constable, and M. M. Chun, “Characterizing Attention with Predictive Network Models,” Trends in Cognitive Sciences, vol. 21, no. 4. Elsevier Ltd, pp. 290–302, 01-Apr-2017.

[13] F. Liem et al., “Predicting brain-age from multimodal imaging data captures cognitive impairment,” Neuroimage, vol. 148, pp. 179–188, Mar. 2017.

[14] S. Gao, A. S. Greene, R. T. Constable, and D. Scheinost, “Combining multiple connectomes improves predictive modeling of phenotypic measures,” Neuroimage, vol. 201, p. 116038, Nov. 2019.

[15] G. Varoquaux, P. R. Raamana, D. A. Engemann, A. Hoyos-Idrobo, Y. Schwartz, and B. Thirion, “Assessing and tuning brain decoders: Cross-validation, caveats, and guidelines,” Neuroimage, vol. 145, pp. 166–179, Jan. 2017.

[16] G. Varoquaux, “Cross-validation failure: Small sample sizes lead to large error bars,” NeuroImage, vol. 180. Academic Press Inc., pp. 68–77, 15-Oct-2018.

[17] M. D. Rosenberg et al., “Functional connectivity predicts changes in attention observed across minutes, days, and months,” Proc. Natl. Acad. Sci. U. S. A., vol. 117, no. 7, pp. 3797–3807, Feb. 2020.

[18] S. W. Yip, D. Scheinost, M. N. Potenza, and K. M. Carroll, “Connectome-based prediction of cocaine abstinence,” Am. J. Psychiatry, vol. 176, no. 2, pp. 156–164, Feb. 2019.

[19] A. S. Greene, S. Gao, D. Scheinost, and R. T. Constable, “Task-induced brain state manipulation improves prediction of individual traits,” Nat. Commun., vol. 9, no. 1, Dec. 2018.

[20] E. M. R. Lake et al., “The Functional Brain Organization of an Individual Allows Prediction of Measures of Social Abilities Transdiagnostically in Autism and Attention-Deficit/Hyperactivity Disorder,” Biol. Psychiatry, vol. 86, no. 4, pp. 315–326, Aug. 2019.

[21] D. Opitz and R. Maclin, “Popular Ensemble Methods: An Empirical Study,” J. Artif. Intell. Res., vol. 11, pp. 169–198, Aug. 1999.

[22] L. Breiman, “Bagging Predictors,” 1996.

[23] J. Richiardi, H. Eryilmaz, S. Schwartz, P. Vuilleumier, and D. Van De Ville, “Decoding brain states from fMRI connectivity graphs,” Neuroimage, vol. 56, no. 2, pp. 616–626, May 2011.

[24] A. Nikolaidis, A. S. Heinsfeld, T. Xu, P. Bellec, J. Vogelstein, and M. Milham, “Bagging improves reproducibility of functional parcellation of the human brain,” Neuroimage, p. 116678, Feb. 2020.

[25] L. Wei, B. Jing, and H. Li, “Bootstrapping promotes the RSFC-behavior associations: An application of individual cognitive traits prediction,” Hum. Brain Mapp., p. hbm.24947, Mar. 2020.

[26] D. C. Van Essen, S. M. Smith, D. M. Barch, T. E. J. Behrens, E. Yacoub, and K. Ugurbil, “The WU-Minn Human Connectome Project: An overview,” Neuroimage, vol. 80, pp. 62–79, Oct. 2013.

[27] M. E. Calkins et al., “The Philadelphia Neurodevelopmental Cohort: Constructing a deep phenotyping collaborative,” J. Child Psychol. Psychiatry Allied Discip., vol. 56, no. 12, pp. 1356–1369, Dec. 2015.

[28] W. B. Bilker, J. A. Hansen, C. M. Brensinger, J. Richard, R. E. Gur, and R. C. Gur, “Development of Abbreviated Nine-Item Forms of the Raven’s Standard Progressive Matrices Test,” Assessment, vol. 19, no. 3, pp. 354–369, 2012.

[29] T. M. Moore, S. P. Reise, R. E. Gur, H. Hakonarson, and R. C. Gur, “Psychometric properties of the penn computerized neurocognitive battery,” Neuropsychology, vol. 29, no. 2, pp. 235–246, Mar. 2015.

[30] R. C. Gur et al., “A cognitive neuroscience-based computerized battery for efficient measurement of individual differences: Standardization and initial construct validation,” J. Neurosci. Methods, vol. 187, no. 2, pp. 254–262, Mar. 2010.

[31] D. C. Van Essen et al., “The Human Connectome Project: A data acquisition perspective,” Neuroimage, vol. 62, no. 4, pp. 2222–2231, Oct. 2012.

[32] T. D. Satterthwaite et al., “Neuroimaging of the Philadelphia Neurodevelopmental Cohort,” NeuroImage, vol. 86. Academic Press Inc., pp. 544–553, 01-Feb-2014.

[33] M. F. Glasser et al., “The minimal preprocessing pipelines for the Human Connectome Project,” Neuroimage, vol. 80, pp. 105–124, Oct. 2013.

[34] A. Joshi et al., “Unified framework for development, deployment and robust testing of neuroimaging algorithms,” Neuroinformatics, vol. 9, no. 1, pp. 69–84, Mar. 2011.

[35] S. M. Smith et al., “Advances in functional and structural MR image analysis and implementation as FSL,” in NeuroImage, 2004, vol. 23, no. SUPPL. 1, pp. S208–S219.

[36] R. Frackowiak et al., Human Brain Function 2nd Edition. Chapter 6: Morphometry. 2003.

[37] X. Shen, F. Tokoglu, X. Papademetris, and R. T. Constable, “Groupwise whole-brain parcellation from resting-state fMRI data for network node identification.,” Neuroimage, vol. 82, pp. 403–15, Nov. 2013.

[38] X. Shen et al., “Using connectome-based predictive modeling to predict individual behavior from brain connectivity,” Nat. Protoc., vol. 12, no. 3, pp. 506–518, Feb. 2017.

[39] W.-T. Hsu, M. D. Rosenberg, D. Scheinost, R. T. Constable, and M. M. Chun, “Resting-state functional connectivity predicts neuroticism and extraversion in novel individuals,” Soc. Cogn. Affect. Neurosci., vol. 13, no. 2, pp. 224–232, Jan. 2018.

[40] J. D. Hunter, “Matplotlib: A 2D Graphics Environment,” Comput. Sci. Eng., vol. 9, no. 3, pp. 90–95, 2007.

[41] M. Waskom et al., “mwaskom/seaborn: v0.8.1 (September 2017).” Sep-2017.

[42] H. Jiang et al., “Predicting Brain Age of Healthy Adults Based on Structural MRI Parcellation Using Convolutional Neural Networks,” Front. Neurol., vol. 10, p. 1346, Jan. 2020.

[43] D. Scheinost et al., “Ten simple rules for predictive modeling of individual differences in neuroimaging,” NeuroImage, vol. 193. Academic Press Inc., pp. 35–45, 01-Jun-2019.

[44] R. A. Poldrack, G. Huckins, and G. Varoquaux, “Establishment of Best Practices for Evidence for Prediction: A Review,” JAMA Psychiatry, vol. 77, no. 5. American Medical Association, pp. 534–540, 01-May-2020.

[45] B. Mwangi, T. S. Tian, and J. C. Soares, “A review of feature reduction techniques in Neuroimaging,” Neuroinformatics, vol. 12, no. 2. Humana Press Inc., pp. 229–244, 2014.

[46] D. S. Barron et al., “Task-Based Functional Connectomes Predict Cognitive Phenotypes Across Psychiatric Disease,” bioRxiv, p. 638825, May 2019.

[47] “Overfitting | Meaning of Overfitting by Lexico.” [Online]. Available: https://www.lexico.com/definition/overfitting. [Accessed: 28-May-2020].

[48] T. D. Satterthwaite et al., “Linked Sex Differences in Cognition and Functional Connectivity in Youth.,” Cereb. Cortex, vol. 25, no. 9, pp. 2383–94, Sep. 2015.

[49] C. Zhang, C. C. Dougherty, S. A. Baum, T. White, and A. M. Michael, “Functional connectivity predicts gender: Evidence for gender differences in resting brain connectivity,” Hum. Brain Mapp., vol. 39, no. 4, pp. 1765–1776, Apr. 2018.

[50] R. F. Betzel, L. Byrge, Y. He, J. Goñi, X. N. Zuo, and O. Sporns, “Changes in structural and functional connectivity among resting-state networks across the human lifespan,” Neuroimage, vol. 102, no. P2, pp. 345–357, Nov. 2014.

[51] N. U. F. Dosenbach et al., “Prediction of individual brain maturity using fMRI,” Science (80-.)., vol. 329, no. 5997, pp. 1358–1361, Sep. 2010.

